# “Technical Note: DeepLabCut-Display: open-source desktop application for visualizing and analyzing two-dimensional locomotor data in livestock”

**DOI:** 10.1101/2023.10.30.564795

**Authors:** Jacob Shirey, Madelyn P. Smythe, L. Savannah Dewberry, Kyle Allen, Eakta Jain, Samantha A. Brooks

## Abstract

**Abstract:** Gait assessments are a key part of determining the wellbeing of livestock. Techniques for gait assessment have traditionally involved human-eye inspections or reflective markers, but markerless computer vision methods have been developed in recent years. Despite many computer vision tools providing high-quality pose estimations in an efficient manner, they lack post-processing functionality. A review of model performance and calculation of gait parameters is a necessary step to fully harness the capability of this new technology. Thus, this study developed DeepLabCut-Display, an open-source desktop software application. DeepLabCut-Display allows a user to upload the video and coordinate data associated with the output of DeepLabCut, a prominent pose-estimation software tool. A user can review the video and coordinate data in parallel, filter points by a likelihood threshold, and automatically calculate gait parameters. Specific video frames, filtered data, and gait parameters can be exported from the application for further usage. The source code is publicly hosted on GitHub alongside installation and usage instructions. DeepLabCut-Display, the product of interdisciplinary and collaborative design between software developers and animal scientists, will alleviate a critical bottleneck in processing of data for locomotor analysis in livestock.

**Summary Statement:**

a. DeepLabCut-Display is a utility to dynamically visualize raw marker coordinates, and to automatically produce gait parameters for locomotion analysis of horses and other livestock.

**Lay Summary:** Artificial intelligence systems that can predict and track the positions of objects are now being applied in many fields, including animal science. Veterinarians and animal scientists use these systems to create pose estimations, a digital label of anatomical landmarks overlaid on a video of an animal in motion. They are used to quantify the subject’s motion and detect anomalies that may be indicative of disease or injury. Pose estimation systems are efficient and accurate, but they lack features like data visualization and post-processing analysis that are necessary to make determinations about the animal’s motion. This study developed DeepLabCut-Display, a software application that can visualize the data from a pose estimation system and provides a set of tools for further analysis. After a user is done with analysis, they can save the results to their computer. The application was made by a collaboration between software developers and animal scientists, highlighting how interdisciplinary teams are effective at producing useful software.

## 6. Introduction

Terrestrial locomotion, or the way in which land animals move, is an incredibly complex behavior. Taking just one single step involves the coordinated motion of many bones, joints, muscles and tendons. This combination of swinging, pushing and pulling results in a non-linear path of motion for any individual point on the animal. Observation of locomotion in livestock species is a key tool for veterinarians and animal scientists, as posture and movement often indicate health status (Becker et al., 2013). Understanding healthy locomotion allows us to better identify unhealthy movement patterns and recognize injury and illness.

Techniques for studying locomotion phenotypes, known more collectively as gait assessments, can involve manual analysis by a human observer (Menzies-Gow et al., 2010), surgically-implanted electromyographic sensors (Lavoie et al., 1995), motion capture systems with reflective markers (Wickler et al., 2004), or instrument packaged with three-dimensional gyroscopes or accelerometers that track the motion and orientation of the animal (McCracken et al., 2012; Fischer et al., 2022). The goal of all of these approaches is to estimate anatomical landmark points, or points of interest on the body, such as the knees, so that the motion of the animal can be quantified.

Yet, many of these traditional techniques have serious drawbacks. Attaching sensors to the animal can be intrusive and alter their natural gait. This is also a time consuming and labor intensive approach in addition to costs incurred to procure sensors and errors introduced when, for example, sensors fall off. Human eye-based observations are not repeatable nor objective (de Mira et al., 2019), and cannot detect or quantify subtle changes. Additionally, livestock may also be kept in large herds making gait analysis of a population expensive and infeasible. Recent advances in deep neural networks enable markerless estimation methods by dramatically increasing the throughput of video labeling. Individual frames of captured video can be annotated for anatomical landmarks with pose estimation via a computer vision algorithm. This not only provides the precision and speed of marker based tracking, but also eliminates the need for invasive physical markers placed on the animal.

One of the leading scientific tools for pose estimation of animals is DeepLabCut (DLC). DLC is an open source Python-based software package that can annotate poses on all animals. DeepLabCut utilizes deep neural networks to analyze images or video frames for pose estimation with great accuracy (Mathis et al., 2018). This tool is used heavily in studying lab rodents, but is able to be applied to all types of animals as a result of transfer learning. DLC is particularly useful in situations where ideal laboratory conditions cannot be easily created. Ecologists conduct field studies by monitoring wildlife with trail cameras. It is impossible to employ traditional marker-technology to study these animals in their natural condition. Wiltshire et al. used DLC to make pose estimates of chimpanzees and bonobos. They found that the machine learning models produced pose estimations with a consistency similar to or better than many different human labeling (Wiltshire et al., 2023), and in a fraction of the time required for human labeling. The model was also able to work in a variety of environments, from open clearings to dense forest.

Large scale farm animal operations also benefit from the portability of DLC processed pose estimation. Broiler chickens can develop lameness that hampers production and animal welfare. Diagnosing lameness requires that subjective human experts score individual chickens by eye. With DLC, researchers were able to score gait by transferring another chicken model (Doornweerd et al., 2021) and adding only 181 additional frames to the dataset (Fodor et al., 2023). Genetic phenotypes can also be quickly observed and analyzed for breeding programs. One study involving pigs was able to track joint locations with an error of only 3.3cm (Gorssen et al., 2022).

DLC, though it performs well for pose estimation, falls short of providing human-readable metrics to the user. DLC outputs include a video file of the original input annotated with landmarks, and a spreadsheet of the landmark coordinates for each frame. While this output is accurate and descriptive, it lacks visualizations useful for a human reviewer to graphically assess model performance, or for generation of quantitative measures of locomotor function. Model revision is difficult without a means to synchronize tabular and video data and identify scenarios where DLC produced an errant prediction, despite a strong log-likelihood score. We sought to create an easy-to-use graphical interface for this process. Additionally, we desired to automate the calculation of gait parameters to the point where physiologists and industry professionals could begin to apply the DLC tool in the field, without extensive knowledge of machine learning and computer science.

Therefore we pursued a new software tool to meet the need for data visualization and gait parameter calculation from outputs generated by DLC. Known as DeepLabCut-Display, the tool seeks to streamline the analysis process for equine scientists and increase the leverage of artificial intelligence applications in the field of animal science.

## 7. Methods

We chose to develop the tool as a standalone desktop application. Software that can run locally on a user’s computer does not require the maintenance of a remote server. This allowed us to focus on the development of the tool’s functionality. Python 3.10 is the sole programming language used to write the application (Python Software Foundation, 2023). Python was chosen for its versatility, supporting many well-maintained open source libraries for data analysis, user-interfaces, and machine learning. Its ease of use, due to its nature as a dynamically typed and interpreted language, made it much quicker to develop a working prototype.

The application must be able to visualize the data output from DeepLabCut. This output comes in comma-separated value (CSV) format. The file’s columns list the horizontal pixel coordinate, vertical pixel coordinate, and prediction likelihood of each landmark point. The rows specify the values for each frame in the analyzed video. We used the Pandas package to manage this coordinate data in the application (McKinney, 2011). This package can read and store tabular data within a Python data structure. It also boasts a large number of functions to manipulate the coordinate data.

DeepLabCut also outputs a copy of the original video input annotated with the landmark points. This video needs to be opened and viewed within the application. We chose to use the OpenCV python package to open the video and extract individual frames (Bradski, 2000).

We needed to plot the coordinate data with respect to each video frame to display how the values changed over time. We used a set of line graphs to display the coordinates for each landmark point. The data plots are rendered using MatPlotLib (John D. Hunter, 2007). This is a 2D graphics package that is widely used for data plotting and visualization.

Our application needed to allow the user to interact with the data. This includes importing the data, browsing through different displays, performing calculations, and exporting results from the application. We created a graphical user interface (GUI) for the application to achieve this goal. It was written using the PyQt framework (Riverbank Computing). This is a Python adaptation of the Qt library for creating a GUI.

Two of the gait parameters were based on calculations in a related paper (Dewberry et al., 2023). These were the stride length and duty factor of each stride. These calculations were originally written in the MATLAB programming language. For our application, the mathematical and algorithmic details were replicated in Python. In addition to the Python packages listed above, the SciPy package was employed to implement signal filtering (Virtanen et al., 2020).

The application source code was developed and tested in a Python virtual environment. Version control throughout development was managed by Git (Spinellis, 2012). The code repository is hosted on GitHub, at https://github.com/jakeshirey/DeepLabCut-Display. The repository includes links to installation and usage video tutorials, in addition to the necessary source code to build the application.

## 8. Results

### a. User Interface

DeepLabCut-Display offers a simple user interface with two primary components: a frame viewer and a set of plotted charts. A set of buttons that initiate functionality runs along the bottom of the window. The components can be adjusted to take up more or less space based on the user’s preference.

The frame viewer allows a user to upload a video file and view individual frames. This component is made in the style of a typical video player, but does not have playback functionality nor display timestamps. Instead, it shows the index of the current frame and only allows for scrubbing through the frames using a slider. This design choice focuses on the purpose of the tool: to identify frames where points may be mislabeled by the model, despite a confident log likelihood score. Users of this application are intended to spend their efforts pausing and analyzing specific frames, rather than watching the full video. If a user wishes to save a specific frame for future reference, the application can save individual frames of the video as an image file.

The plotted charts visualize the coordinate data given as an output by DeepLabCut. Upon uploading the coordinate data file into the application, each of the landmark point names are loaded into a list visible on the right side of the charts. Clicking on one of the landmark names plots the associated data onto each of the charts. The two charts are labeled “Pixel Coordinate (X)” and “Pixel Coordinate (Y)”: both the horizontal and vertical video coordinates are plotted separately to distinguish the motion by dimension. The coordinates are plotted with respect to the video frames to highlight the change in value over time. The name of each landmark point is displayed in a list on the side of the charts. Clicking on an item in the list plots the coordinate data for that landmark. Selective plotting allows a user to see only the data that they may need for analysis.

Synchronization between the two data modalities is a key element in this tool. For a user to carefully assess and interpret the coordinates, it is imperative to see the corresponding video frames. DeepLabCut-Display matches the state of the frame viewer to the state of the charts. A bright red vertical line is drawn through each of the plots intersecting the point representing the current rendered video frame. When the video frame changes, the vertical line moves. Users can also click on the graph to summon the vertical line to that point, and the video will switch to that frame.

### b. Data Transformation

Feedback from animal scientist users revealed a need to display only data of certain characteristics on the plots. For every coordinate point, DeepLabCut produces an associated likelihood value. This value indicates how confident the algorithm is that the labeled point is the true landmark, with values closer to one being more confident.

DeepLabCut-Display supports filtering points by likelihood. A button below the plots opens a pop-up window that queries the user for a numerical input between zero and one. When applied, the plot will only display coordinate data with an associated likelihood value equal to or above the input value. All other points are removed from the plot. A user can export the filtered data from the application and the values that were filtered out are replaced by null values. How this null value is represented depends on the computer’s operating system, version of Python in the application, and the program in which the transformed values are viewed. This may be useful for post-export statistical analyses on platforms that allow for null values to be ignored.

### c. Gait Parameter Calculations

The loaded coordinate data can be used to calculate gait parameters from within the application. We included six gait parameters based on commonly used measures in livestock and rodent gait analysis research (Wickler et al., 2006; Dewberry et al., 2023). These are listed in table 1, with descriptions and formulas for calculation.

**Table 1.**
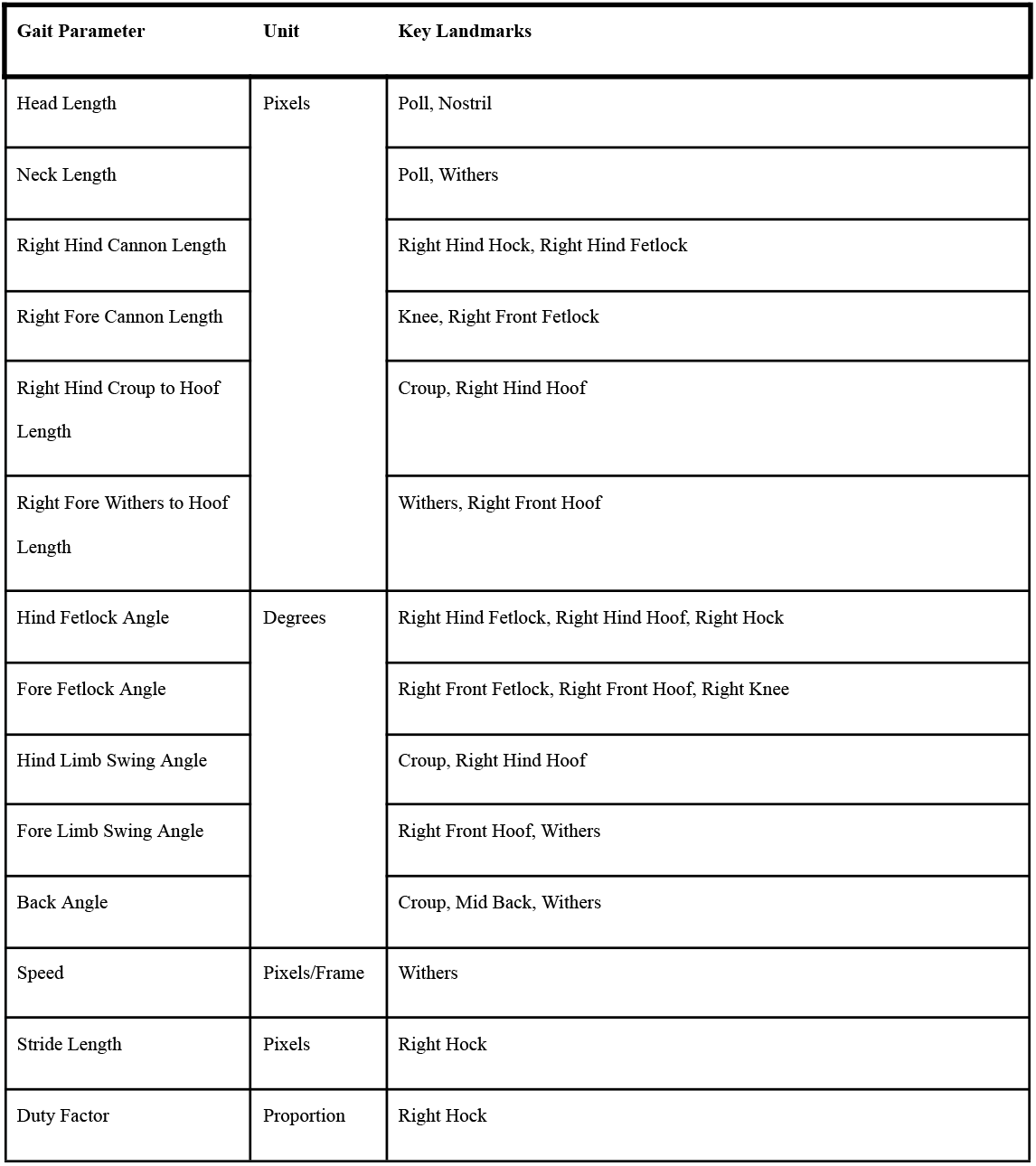
The supported gait parameters in DeepLabCut-Display.

Additionally, a number of linear conformational measures between landmarks on the body of the animal that provide additional utility in locomotion assessment. Our tool was built with initial application in the equine sciences in mind, so the measures were piloted using data for the horse.

On the main application page, a button labeled “Calculate Gait Parameters” initiates the calculation process. When clicked, a pop-up window is displayed to the user asking which gait parameters to be calculated, and which summary statistics to include. These summary statistics are a value calculated over the entire set of frames for each parameter, and include the minimum, maximum, mean, and standard deviation. When the user is satisfied with their selections, they select a button within the pop-up window labeled “Calculate”. The parameters and summary statistics are generated, and the user is then prompted to save the results into a spreadsheet-like file.

## 9. Discussion

### a. Advantages

DeepLabCut-Display allows for animal scientists to harness the full power of DeepLabCut. While the neural network model behind DeepLabCut is powerful, the program lacks in utilities for visualizing and understanding the model’s predictions. Our program provides an accessible means of initially evaluating the DeepLabCut output. The user interface design is minimal to reduce the learning curve of the application.

This tool was created by a team of software developers, who worked in conjunction with animal scientists to translate stakeholder needs into concrete functionality and features. Development in an interdisciplinary environment was highly productive as it allowed for a synthesis of techniques known by animal scientists to be translated into programmatic logic written by a software developer.

### b. Disadvantages

This program initially has a focus specific to horses (*Equus caballus*) as video data collection is easily accomplished in this species, and the horse industry has a number of immediate applications for gait analysis. The gait parameter calculators included in the analysis part of the toolkit are exclusively for use with the set of landmarks initially defined for the body of the horse. However, additional pre-trained models are in development for several livestock species using these same landmarks, and our goal is to ultimately create a suite of tools enabling adaptation of this workflow to diverse agricultural species.

It is important to note that although the gait parameters are custom built for horses, DeepLabCut-Display is able to read and render any DeepLabCut output file. Any landmark data and corresponding video can be viewed in DeepLabCut-Display, and many of the gait parameters, although designed for horses, could be usefully applied to other quadrupedal species.

### c. Limitations

DeepLabCut-Display is hosted as a set of source code files on a GitHub repository, and not as a stand-alone executable file. The user must go through the process of downloading, compiling, and installing the files in order to use the program. This limitation is due to the underlying operating system of the user: different machines have different requirements. To overcome this, we have created a written and video guide to accompany the program. These guides are linked on the program’s GitHub page.

## 10. Conclusion

DeepLabCut-Display is an open-source tool for landmark data visualization and gait parameter calculation. It fills a procedural gap in computer-vision based gait analysis. This tool makes review of model performance easier for animal scientists who may not have software development skills. Currently, the source code of the application is hosted at https://github.com/jakeshirey/DeepLabCut-Display with download and installation instructions. We hope to continue to expand on the processing capabilities of this tool by adding parameters for more species and new robust analysis features.

## 11. Acknowledgments

Our sincere thanks to the many undergraduate researchers working on the livestock locomotion project.

## 14. Data availability

**https://github.com/jakeshirey/DeepLabCut-Display**

This is the link to the GitHub repository hosting the source code of the application.

## 16. Figure Legends

**Figure 1.**
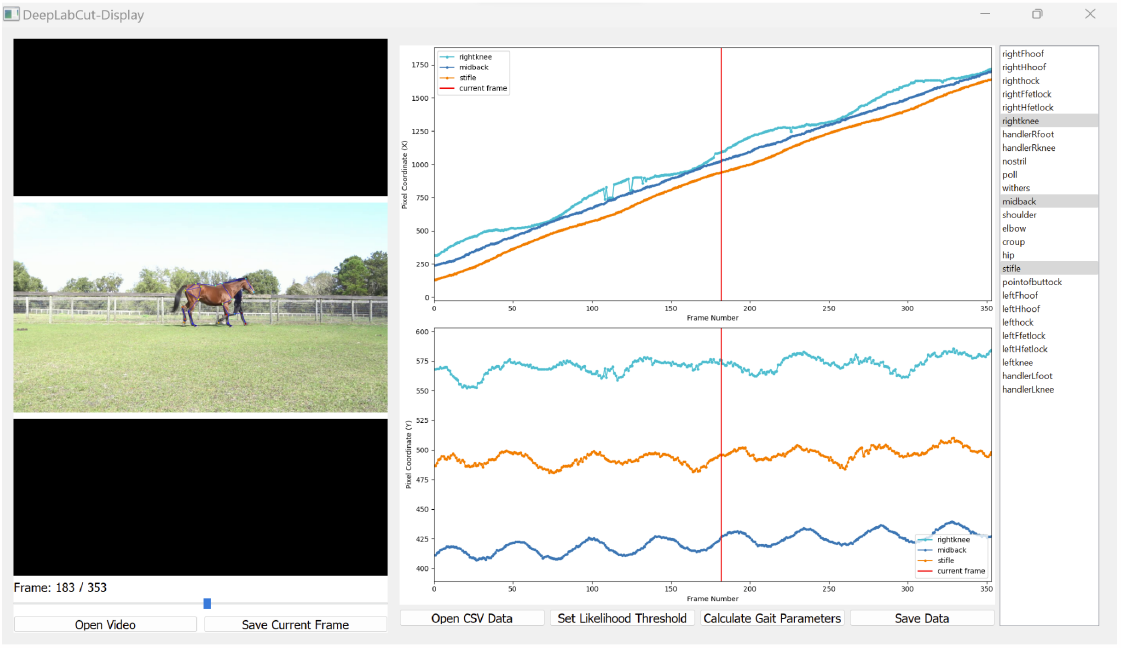
DeepLabCut-Display user interface. The application is split into two main panels, a video frame display and coordinate data plot. The coordinate data plot shows the horizontal coordinate (top) and vertical coordinate (bottom) in separate axes with respect to time The vertical red line in the coordinate data plot denotes the exact frame currently being displayed. The three highlighted landmarks (rightknee midback, stifle) on the right column are the three lines being displayed on the plots.

## 17. Appendices

Supplemental File 1 contains a description of the equations used to calculate parameters in the DeepLabCut-Display tool.

