## Supplemental File 1 for "“Technical Note: DeepLabCut-Display: open-source desktop application for visualizing and analyzing two-dimensional locomotor data in livestock”"

**Equation 1.** The joint angle formula implemented in DeepLabCut-Display.

$$angle = \tan^{-1}\left(\frac{det\left(\begin{bmatrix} \mathbf{a} \\ \mathbf{b} \end{bmatrix}\right)}{\mathbf{a} \cdot \mathbf{b}}\right)$$

where

$$\mathbf{a} = \begin{bmatrix} e_{1_x} - v_x & e_{1_y} - v_y \end{bmatrix}$$

and

$$\mathbf{b} = \begin{bmatrix} e_{2_x} - v_x & e_{2_y} - v_y \end{bmatrix}$$

**a** and **b** are the row vectors representing the horizontal and vertical distances between the vertex and each of the endpoints respectively.
